# Identifying Restrictions in the Order of Accumulation of Mutations during Tumor Progression: Effects of Passengers, Evolutionary Models, and Sampling

**DOI:** 10.1101/005587

**Authors:** Ramon Diaz-Uriarte

**Affiliations:** Dept. Biochemistry, Universidad Autónoma de Madrid Instituto de Investigaciones Biomédicas “Alberto Sols” (UAM-CSIC) Madrid, Spain

## Abstract

Cancer progression is caused by the sequential accumulation of mutations, but not all orders of accumulation of mutations are equally likely. When the fixation of some mutations depends on the presence of previous ones, identifying restrictions in the order of accumulation of mutations can lead to the discovery of therapeutic targets and diagnostic markers. Using simulated data sets, I conducted a comprehensive comparison of the performance of all available methods to identify these restrictions from cross-sectional data. In contrast to previous work, I embedded restrictions within evolutionary models of tumor progression that included passengers (mutations not responsible for the development of cancer, known to be very common). This allowed me to asses the effects of having to filter out passengers, of sampling schemes, and of deviations from order restrictions. Poor choices of method, filtering, and sampling lead to large errors in all performance metrics. Having to filter passengers lead to decreased performance, especially because true restrictions were missed. Overall, the best method for identifying order restrictions were Oncogenetic Trees, a fast and easy to use method that, although unable to recover dependencies of mutations on more than one mutation, showed good performance in most scenarios, superior to Conjunctive Bayesian Networks and Progression Networks. Single cell sampling provided no advantage, but sampling in the final stages of the disease vs. sampling at different stages had severe effects. Evolutionary model and deviations from order restrictions had major, and sometimes counterintuitive, interactions with other factors that affected performance. This paper provides practical recommendations for using these methods with experimental data. Moreover, it shows that it is both possible and necessary to embed assumptions about order restrictions and the nature of driver status within evolutionary models of cancer progression to evaluate the performance of inferential approaches.

## Introduction

Cancer progression is caused by the sequential accumulation of somatic mutations, including changes in copy number (structural variants), single nucleotides (SNP variants) and DNA methylation patterns during the life of an individual [1–3]. Among the mutations causally responsible for the development of cancer (*drivers*) not all possible orders of accumulation seem equally likely, and the fixation of some mutations can depend on the presence of other mutations. For example, in colorectal cancer APC mutations are an early event that precedes mutations in KRAS [4–6]. Understanding the restrictions in the temporal order of accumulation of driver mutations not only provides insights into cancer biology, but can help identify early markers of disease as well as therapeutic targets [5–9] and can be an instrumental tool in the search for the “Achilles’ Heel” of oncogene addiction [3, 10, 11]. In addition, understanding the correct order of events is necessary for the assessment of the validity of the genetic context of cell lines and animal models of human cancer [7, 8].

In this context, a variety of methods have been developed to try to infer the possible restrictions in the order of accumulation of driver mutations from cross-sectional data. Longitudinal data would be better suited for this problem but it is much harder to obtain and cross-sectional data is (and will remain) the main source of data (e.g., the growing number of genomes available through international sequencing projects) for addressing these and similar problems [5, 12]. I provide next a brief review of the main methods, including recent developments, but see more extensive reviews in [13, 14]. The oncogenetic tree (OT) model [15] was introduced as an extension of the linear path model [16]: in OTs progression starts from a common (non-altered) root, and branches out, so that there are several mutational pathways that can be observed simultaneously. OTs, by virtue of being trees, can only model order restrictions where an event depends on its single parent. Another early model are distance-based trees [17,18], but their meaning is rather different, since the observed mutations are only placed in the leaves or terminal nodes of the tree, and the internal nodes are unobserved and unknown events, which precludes an interpretation in terms of order restrictions like “mutation A is required for mutation B”. Distance-based trees and other models [19] that do not try to infer order restrictions will not be considered further in this paper.

Conjunctive Bayesian Networks (CBNs) [20] were developed as a generalization of OTs: these are graphs where the occurrence of a mutation can depend on the occurrence of two or more parents (i.e., a *conjunction*). The disease progression models of OTs and CBNs assume that a mutation can only occur with non-negligible probability if the preceding parent mutation(s) in the graph have occurred, which has been called *monotonicity* [12]. Thus, for driver genes, under strict OT and CBN models it would be impossible to observe a genotype that is not compatible with the relations specified in the graph. Less restrictive models for tumor progression were suggested early on, including general Markov models and Bayesian Networks which allow for mutations to occur even if no other aberrations have occurred [14, 21–23]. Progression Networks [12] have been proposed for learning models that include OTs, CBNs, as well as several other special types of Bayesian Networks, and can explicitly incorporate deviations from monotonicity. RESIC [7, 8] differs from other methods because it attempts to find the order of events taking into account the evolutionary dynamics of mutation accumulation. Some recent work [5,22] simultaneously tries to find modules or pathways and their order restrictions; but note that CBNs, OTs, and Progression Networks can be directly applied to module/pathway data, provided those data are partitioned into predefined pathways before the analysis (e.g., [6, 7]).

Having a single graph means having a single set of restrictions that is common to all individuals, but that does not mean that all cells follow the same path (so the actual genotypes and their paths can be quite diverse under one graph). Mixtures of OTs [24] and mixtures of Hidden-variable OTs [25] are one further generalization of OTs where disease progression is modeled allowing for different order restrictions in different subsets of individuals, each one modeled as a (Hidden-variable) OT. By using a star as one of the trees in the mixture, these models can also account for any mutation occurring without its parent(s) having occurred. In this paper I restrict attention to finding a single graph, the approach most widely used in the literature, and one which seems to work even when there are mild departures from the single-graph assumption [26] (but see [27] and Discussion).

Most of the above are general methods, and can be applied to different kinds of data including cytogenetic, gene mutation, and pathway alteration data [6, 15]. This versatility, coupled with the increasing wealth of cross-sectional data available, provides an excellent opportunity to try to understand the still largely unknown details of the order of mutations. However, in spite of the relevance of the problem for both diagnostic and therapeutic purposes, there are very few systematic comparisons of method performance, and they do not provide a clear and robust answer to the question of method choice.

Applied usage of the above methods faces at least three additional major problems. First, most of the mutations present in cancer cells are not *driver* mutations, but *passenger* mutations not responsible for the development of cancer [28–31]. Passenger mutations can show a non-negligible frequency because they “hitchhike” on drivers [1,32]. Unless we know what mutations are drivers, the presence of passengers in our data sets forces us to use some filtering procedure to select which mutations (or, generally, alterations) to use with (or to pass on to) the methods to infer order restrictions. However, the simulations in the only comparison of methods available [13], as well as in the original descriptions of new methods [12, 33, 34], have all been conducted assuming that the identity of the driver mutations is known. Virtually all papers that try to infer order restrictions, including methodological papers, rely on simple frequency-based selection or filtering procedures to select which genes to use [6, 15, 18, 22, 34–38], but the effects of these filtering approaches on the performance of the methods to infer order restrictions are completely unknown.

Second, attention to sampling decisions is largely missing from the literature. OTs, CBNs, and Progression Networks are generative models, and simulations that examine method performance [12, 13, 33, 34] obtain genotypes directly from these generative models. But, except when we use single cell sampling, our experimental data are from samples that aggregate over many cells and the joint and marginal frequencies of mutations of those aggregates can depend not only on the aggregation *per se* but also on when we sample (due, for example, to the clonal expansion episodes), and differ greatly from distributions obtained from the generative models.

Finally, development and evaluation of methods of reconstruction of order restrictions are conducted without consideration for the evolutionary model of tumor progression (but see [7, 8] and Discussion). This problem is highlighted by Sprouffske et al. [39]: referring to oncogenetic tree models they say (p. 1136) “This is not an evolutionary model because the oncogenetic tree does not represent ancestral relationships within a neoplasm but rather a summary of the observed co-occurrences of mutations across independent neoplasms”. This lack of consideration for the evolutionary model is also unfortunate since it does not provide a clear mechanistic interpretation of (nor a simple mechanistically-based procedure for generating) deviations from the restrictions encoded in the graph. Of particular interest is monotonicity (a mutation in a driver gene can only be observed if the preceding parent mutations in the graph have occurred), because deviations from it can easily arise when a mutation behaves as a driver or as a passenger depending on the genetic context—i.e., depending on which other genes are mutated [32, 40].

As we have seen, data simulated from the generative models of OTs, CBNs and Progression Networks cannot be used to address any of those three problems (passengers, sampling, deviations from monotonicity). However, it is possible to incorporate the order restrictions encoded in CBNs, OTs, and Progression Networks into plausible evolutionary models of tumor progression (in fact, recently a simulation tool that incorporates simple order restrictions among four drivers has been published [41] —see Discussion). If we model together drivers (with possible restrictions) and passengers we can address the consequences of having to filter drivers from passengers. Incorporating order restrictions within evolutionary models would also allow us to address two questions of immediate practical relevance related to data collection: should we try to use single cell sampling now that it is becoming a realistic possibility [42] instead of whole tumor sampling? and would it be better to try to use samples collected in the final stages of the disease vs. using samples collected also at intermediate stages? Finally, using explicit evolutionary tumor growth models also allows us to examine the consequences of deviations from monotonicity and the genetic context dependence of driver status. In fact, we can generate data using simulations in a way that closely mimics the process of data generation and order restriction inference from patient data, as illustrated in Figure 1.

**Figure 1.**
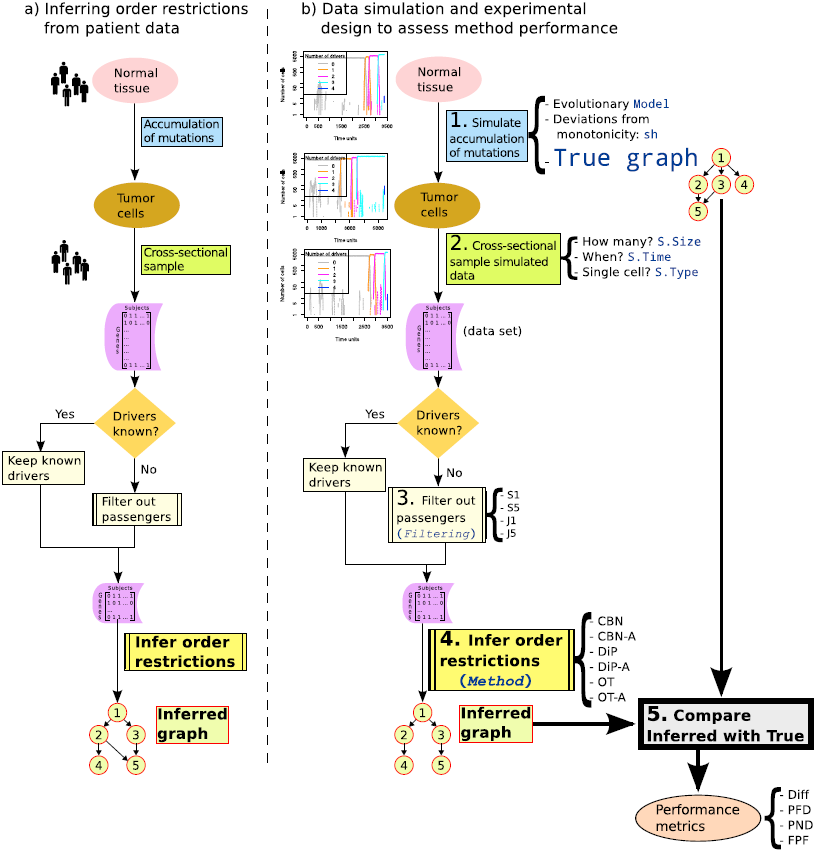
Inferring order restrictions. (a) Main steps in the analysis of patient data. (b) Main steps used in this paper for the generation (simulation) of data and its analysis. Terms in monospaced blue font are those in Table 1, and terms in italics, as in Table 1, correspond to within-data set factors. Numbers indicate the chronological order of the steps. In step 1, cancer development is simulated for the specified values of Model, sh, and True Graph. This simulation generates tumor cell data for the equivalent of a single patient in panel (a). In step 2, data for S.Size patients are sampled (cross-sectional sampling) according to the settings of S.Time and S.Type, producing a data set (a collection of genotypes: a matrix of subjects by genes). If the identity of the true drivers is not known, Filtering in step 3 removes from the data set the genes that do not meet certain frequency criteria. The data set is then passed on, in step 4, to one of the specified methods to infer the graph that encodes the order restrictions. This inferred graph is compared, in step 5, with the true graph (which was used in step 1 to generate the cancer cell data) yielding the four performance metrics Diff, PFD, PND and FPF. The process illustrated here was repeated 20 times for all possible combinations of Model, sh, True Graph, S.Time, S.Type, S.Size. Every data set was subject to all Filtering procedures and analyzed with all six Methods.

In this paper I incorporate the order restrictions into evolutionary models to address how the performance of all available methods for inferring order restrictions is affected by: a) passenger mutations that lead to uncertainty about the identity of the true drivers and the need to use filtering approaches; b) sampling choices (when and how and how many to sample); c) type of underlying true graph, including presence/absence of conjunctions; d) deviations from the order restrictions encoded in the graphs (deviations from monotonicity); e) evolutionary model of tumor progression.

## Materials and Methods

Details of each factor examined are discussed below, but Table 1 provides an overview of the main factors, and Figure 1 a schema of all the steps. We will deal separately with two different scenarios: one where we know the true identity of the drivers, which we will call “Drivers Known”, and another scenario that replicates the common situation where the data includes both passengers and drivers and we do not know with certainty which is which, a scenario we will refer to as “Drivers Unknown”. The “Filtering” factor is only relevant for the “Drivers Unknown” scenario (as shown in Figure 1 by the “No” path from the decision diamond “Drivers known?”). “Graph”, refers to the structure that encodes the order restrictions and I will reserve the term “tree” for graphs that have no conjunctions.

**Table 1.**
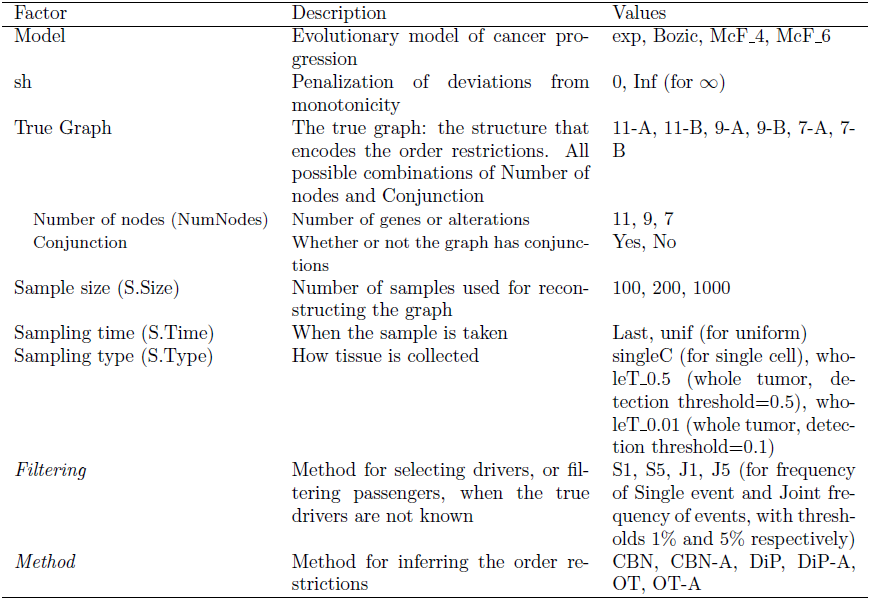
Factors considered and their levels or possible values, together with the acronyms used through the text. The within-data set factors, Filtering and Method (see text), are shown in italics. All other factors are among-data set factors. Sampling scheme, used through the text, refers to when (S.Time) and how (S.Type) we sample.

| Factor | Description | Values |
| --- | --- | --- |
| Model | Evolutionary model of cancer progression | exp, Bozic, McF_4, McF_6 |
| *sh* | Penalization of deviations from monotonicity | 0, Inf (for ∞) |
| True Graph | The true graph: the structure that encodes the order restrictions. All possible combinations of Number of nodes and Conjunction | 11-A, 11-B, 9-A, 9-B, 7-A, 7-B |
| Number of nodes (NumNodes) | Number of genes or alterations | 11, 9, 7 |
| Conjunction | Whether or not the graph has conjunctions | Yes, No |
| Sample size (S.Size) | Number of samples used for reconstructing the graph | 100, 200, 1000 |
| Sampling time (S.Time) | When the sample is taken | Last, unif (for uniform) |
| Sampling type (S.Type) | How tissue is collected | singleC (for single cell), wholeT_0.5 (whole tumor, detection threshold=0.5), wholeT_0.01 (whole tumor, detection threshold=0.1) |
| *Filtering* | Method for selecting drivers, or filtering passengers, when the true drivers are not known | S1, S5, J1, J5 (for frequency of Single event and Joint frequency of events, with thresholds 1% and 5% respectively) |
| *Method* | Method for inferring the order restrictions | CBN, CBN-A, DiP, DiP-A, OT, OT-A |

Method (Drivers Known scenario) or Filtering by Method combinations (Drivers Unknown scenario) were applied to a data set: a matrix of individuals or subjects by genes. Each simulation run, when we sample from it, produces the observed genotype of a subject, and a data set is made from the genotypes of multiple subjects (that are sampled in the same way). The same data set can be analyzed with different Methods (or different Filtering by Method combinations) but for other factors (e.g., Model) different settings of the factor (and reruns of the same settings with different seeds of the random number generator) generate different data sets. This allows us to use an experimental design with among-and within-data set factors so as to examine the effect of Method and Filtering, controlling for possible among-data set variation (as explained below), and so as to minimize computational costs. For factors Model, sh, True Graph (= Number of Nodes * Conjunction), S.Size, S.Type, and S.Time, the among-data set factors, I used a full factorial design (thus, 4 ∗ 2 ∗ 3 ∗ 2 ∗ 3 ∗ 3 ∗ 2 = 864 among-data set factor combinations). For every combination of the among-data set factors I used twenty independent replicate data sets. Each of the twenty replicate data sets was analyzed with every Method or every Filtering by Method combination (the within-data set factors) to infer a graph from the data (i.e., to try to infer the order restrictions among events). Therefore, a total of 864 ∗ 6 = 5184 or 864 ∗ 6 ∗ 4 = 20736 factor combinations for the Drivers Known and Drivers Unknown scenarios, respectively, were examined. Every data set was obtained by independently sampling different (and independently generated) simulated tumor trajectories, according to the settings of the among-data set factors. For instance, if sample size was set to 1000, I simulated 1000 tumor progression trajectories as specified by the settings of Model, sh, and True Graph, and sampled each of those trajectories as specified by the settings of S.Time and S.Type (see Figure 1). This process was repeated at every one of the 864 among-data set factors (and, thus, the results shown here are based on 7.49 million simulated trajectories = 4 ∗ 2 ∗ 3 ∗ 2 ∗ 3 ∗ 2 ∗ 20 ∗ (1000 + 200 + 100)).

### Evolutionary models and simulation

Simulated data were generated using different models of tumor progression. The purpose of using several models is not to compare models of tumor development, but to use a range of plausible ones where we embed the order restrictions so that we can examine how the true underlying model could impact the inference of restrictions. Two of the models used, called here “Bozic” (as it is based on [43]) and “exp” have no density dependence and lead to exponential growth. The second set of models, called “McF_4” and “McF_6”, are based on McFarland et al’s work [44] and lead to logistic-like behavior, as death rate depends on total population size. Table 2 summarizes the main parameters used for the tumor progression models. To simulate data, I used the BNB fast stochastic algorithm [45]. For the McFarland model [44], where death rate is density dependent, BNB does not provide an exact simulation, but can be used to provide a very accurate approximation [45]. Details about the models, choice of model parameters, usage of BNB, and examples of simulated trajectories are provided in *Supplementary Material Section* 2.

**Table 2.** Main parameters for each of the tumor progression models. *j* is the number of drivers with their dependencies met, and *p* the number of drivers with dependencies not met. In all cases *s* = 0.1. *s_h_* is set to either 0 (so it has no effect) or *∞* (so fitness of that clone is 0). *N*: population size. *K* = 2000. ^+^: Strictly, birth rate = max(0, (1 + *s*)*^j^*(1 *-s_h_*)*^p^*).

| Model | Birth rate | Death rate | Mutation rate (per gene per unit time) | Cancer reached if |
| --- | --- | --- | --- | --- |
| Bozic | 1 | $(1-s)^j(1+s_h)^p$ | $10^{-5}$ | $> 10^9$ cells |
| exp | $(1+s)^j(1-s_h)^p$ (+) | 1 | $b_j * 10^{-7}$ | $> 10^9$ cells |
| McF_4 | $(1+s)^j/(1+s_h)^p$ | $\log(1+N/K)$ | $5 * 10^{-7}$ | Number of drivers $\geq 4$ |
| McF_6 | $(1+s)^j/(1+s_h)^p$ | $\log(1+N/K)$ | $5 * 10^{-7}$ | Number of drivers $\geq 6$ |

#### Deviations from monotonicity and genetic context dependence of driver status: *sh*

Genetic context dependence of driver/passenger status [32, 40] and deviations from monotonicity (i.e., from the order of events implied by the graph of the oncogenetic model) can be closely related, and affect the performance of methods to infer order restrictions. A mutation in a driver gene for which all the preceding required mutations have occurred (i.e., a mutation in a gene that has its dependencies satisfied) will lead to an increase in fitness (through its increase of the value *j*, for number of drivers, as shown in Table 2). What about mutations in driver genes that do not have their dependencies satisfied? We can formulate the problem in a general context by using the terminology in [46]. Enforcing monotonicity is equivalent to considering such a mutation as a mutation in an essential housekeeping gene, which can be modeled setting *s_h_*, in the notation of [46], to ∞ (so fitness of such clones is zero). Deviations from monotonicity can arise, however, if such mutation is similar to a passenger mutation: it confers no fitness benefits (and if it has no deleterious effect *s_h_* = 0, similar to setting *s_p_* = 0 in [44]). Of course, in none of these two cases (restrictions not satisfied) would the value of *j* be increased (because a driver only increases fitness if its dependencies are satisfied). In the simulations reported here I considered two extreme scenarios: a) no deviations from the graph of the oncogenetic model are allowed, which I will refer to as “sh = Inf” (from ∞); b) drivers without dependencies satisfied are equivalent to passengers with no deleterious effects, which I will refer to as “sh = 0”. Note that the implementations I used to infer CBN and OT incorporate errors [6, 33, 34] and the OT model explicitly allows for errors due to the occurrence of genetic events outside the model implied by the graph of the oncogenetic model [47]. DiP, the method to infer Progression Networks models [12], explicitly incorporates deviations from monotonicity with the parameter *ε*.

### True graphs, number of drivers, and number of passengers

Six different true graphs have been used, three of them with conjunctions (i.e., graphs that could only be perfectly inferred with either CBN or Progression Networks) and three of them without conjunctions (i.e., trees that could be perfectly reconstructed by all methods compared). The trees (the graphs without conjunctions) are derived from the graphs with conjunctions by removing conjunctions. The graphs have 7, 9, and 11 nodes. The number of nodes of the graphs targets the range of nodes commonly considered in studies that try to reconstruct graphs from real cancer data: 10, 11, and 12 in [33], 7 to 11 (including gene and core pathways) in [6], 7 in [15], 11 and 12 in [35], 8 in [26], 7 (modules) in [22], 9 in [48], 17 in [9] and [23], 12 in [49], 6 and 13 in [21], 12 in [14]. The size of graphs was limited to 11 because CBN cannot deal with more than 14 events or nodes and we need to allow for the possible selection of more than 11 events when Drivers are Unknown. The graphs are shown in *Supplementary Material Section* 3, and we will refer to them by the number of events, using post-fix A for conjunction and B for no conjunction. Graph 11-A is the same as Poset 2 in [33] (their Figure 2A), and Graph 7-A is the same as the estimated graph for pancreatic cancer in [6] (their Figure 2B). Graph 9 was created so as to contain an intermediate number of both nodes and conjunctions between the previous two graphs. In addition to the number of nodes and presence/absence and number of conjunctions, the six graphs used differ in other features (such as depth, total number of edges, existence or not of isolated nodes or subgraphs, existence or not of a single parental node, indegree of conjunctions, and outdegree).

**Figure 2.**
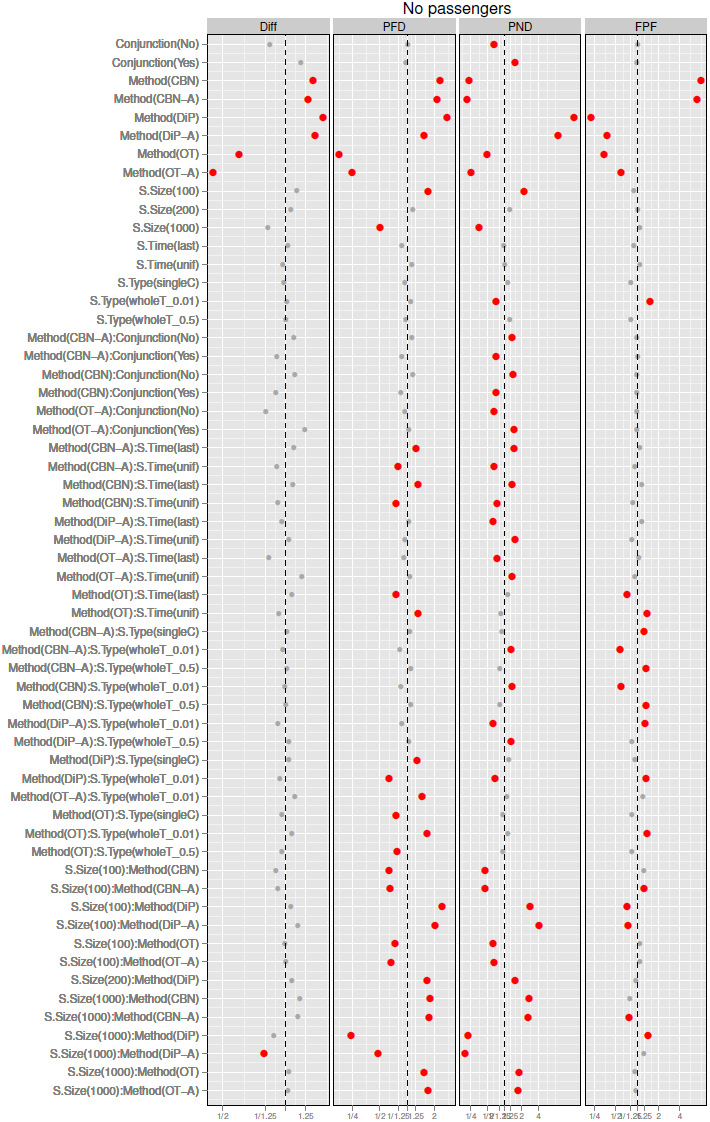
Drivers Known, plot of the coefficients (posterior mean and 0.025 and 0.975 quantiles) for Conjunction, Method, S.Time, S.Type and S.Size from the GLMMs for each of the metrics. X-axis labeled by the exponential of the coefficient (i.e., relative change in the odds ratio or in the scale of the Poisson parameter for Diff): smaller (or lefter) is better. The vertical dashed line denotes no change relative to the overall mean (the intercept). The x-axis has been scaled to make it symmetric (e.g., a ratio of 1.25 is the same distance from the vertical line as a ratio of 1/1.25). Coefficients that correspond to a change larger than 25% (i.e., *ratio >* 1.25 or *<* 1*/*1.25) shown in larger red dots. The coefficients shown are only those that represent a change larger than 25% for at least one metric, or coefficients that are marginal to those shown (e.g., any main effect from an interaction that includes it).

Simulations that used graphs with 11, 9, and 7 seven drivers generated clones with between two and six drivers (see also *Supplementary Material Section* 2), a range which is well within the range of drivers considered in the literature: although some authors [46,50] examine scenarios with 20 drivers, most studies deal with much smaller numbers of drivers [39, 43, 44] and recent reviews suggest that the number of drivers in the cells of most tumors lies between two and six [28, 44]. Regarding number of passengers, it is now widely accepted that most mutations in cancer cells are passengers [28–31, 43, 51]. In the Drivers Unknown scenario I set the proportion of passengers to drivers constant, so that there are four passengers for every driver, a range within that seen in the literature. Our scenario is also relevant if the actual number or fraction of passengers is much larger, but many of those passengers can be excluded *a priori* based on other information, so that they are never considered as candidates for the process of filtering data and inferring graphs (i.e., they are never passed on to step 3 in Figure 1).

### Sample size and sampling type and time

**Sample size (S.Size)** was set to three possible values: 100, 200, 1000. The values considered in other studies, both empirical and methodological, vary widely. For example, 100, 400, and 800 in [35]; 83 to 95, plus a pool of 268 in [6]; 50, 100, 200, 500, and 1000 in [13]; 971 in [26]; 887 in [9]. The set of 100, 200, and 1000 covers a realistic range of sample sizes and will allow us to compare the effects of sample size with those of other factors. **Sampling time (S.Time)** refers to when sampling occurs. S.Time = last means that samples were collected at the end of the simulation (at the end of cancer progression). S.Time = unif (for uniform) means that sampling time was uniformly distributed between the time of appearance of the first mutated driver and the end of the simulation. Uniform sampling is a very simple model for obtaining cross-sectional samples of patients at different stages of the disease. Sampling uniformly between the time of appearance of the first mutated driver and the final stages of the disease is, of course, unrealistic, but I used this type of sampling because it provides a stark contrast with sampling at the end of the disease: S.Time = unif and S.Time = last can be regarded as two extremes of sampling tissue that harbors at least one mutated driver (i.e., cancerous or pre-cancerous tissue). **Sampling type (S.Type)** refers to whether single cell or whole tumor sampling is used, and three values have been used for this factor. When S.Type = singleC (single cell), a simulation provides the genotype of one single cell (or, equivalently, one single clone), where the probability of selecting a clone is proportional to its abundance. When using whole tumor sampling, and as in [39], a biopsy is the entire tumor, but whether a gene is considered mutated or not depends on the detection threshold, and here I used two levels: 0.5 (like [39]) and 0.01, meaning that a gene is considered mutated if it is mutated in 50% or 1%, respectively, of the cells. Of course, it is unlikely that any study using single cell would take a single sample from a patient; however, in this study we focus on cross-sectional data, and single cell sampling is the type of sampling that leads to data most similar to the type of data obtained when we simulate using the generative model of the underlying graph. Moreover, single cell sampling and whole tumor sampling, as used here, can be considered two extremes in the range of sampling possibilities. Likewise, a detection threshold of 0.01 is probably unrealistically low but that setting is used because it combines the capacity of detecting very low frequency co-occurring events (as in single cell sampling) with summing over distinct cells (which could lead to problems similar to the ecological fallacy). The sampling schemes used here ignore any possible spatial structure and tissue architecture [52, 53], not because they are considered irrelevant, but because none of the evolutionary models considered here incorporate them (but note that the uniform sampling scheme can sometimes be equivalent to incorporating spatial structure, if that spatial structure is correlated with time).

### Filtering

When the identity of the driver genes is not known, it is often necessary to select genes before trying to infer the order restrictions. Some studies that deal with chromosomal abnormalities have used the methods of Brodeur and collaborators or Taetle and collaborators, to try to locate non-random breakpoints (see discussion and references in [14, 21]) but these methods are not directly applicable to other types of data. Other authors that deal with chromosomal abnormalities, or that use mutation data, have used one of the following general approaches to decide which alterations to analyze: a) selecting the most frequent mutations, either by setting a minimal number as in [34], [36], and [22], where the seven, 13, or 25, respectively, most frequent alterations are used, or setting a minimal frequency such as in [6], [37], [38], [18] where the threshold is set at 5%, 5%, 10%, 10%, respectively; b) selecting the largest set of events so that every pair of events is observed at least *k* times, as in [47] and [15] where the threshold is five times, out of 124 and 117 cases, respectively.

The key difference between these two filtering procedures, therefore, is that the second uses the joint occurrences of pairs. To comprehensively incorporate common uses, I used four filtering procedures: two of them only consider the marginal frequency of each single event, and I use an “S” to denote “frequency of Single event”, and the other two take into account joint occurrences, and a I use “J” to denote “Joint frequency of pairs of events”. The procedures are S1, that selects any mutation with a frequency larger than 1%, S5 where the threshold is 5%, J1 that selects the largest set of events so that every pair of events is observed at least in 1% of the cases and J5, where the threshold is 5%. In the rare case where a filtering procedure returned more than 12 mutations, the 12 most common were selected.

### Inferring order restrictions: CBN, DiP, OT

I have used three types of methods to infer order restrictions from data: methods that infer OTs, methods that infer CBNs (which should also be able to reconstruct OTs), and methods that infer Progression Networks (and, thus, should be able to reconstruct OTs and CBNs). Each method, when applied to a data set, returns what we will refer to as an “inferred graph” (see Figure 1). For OTs I used the R package Oncotree [54] with its default settings. Some of the analysis were rerun (see Discussion and *Supplementary Material Section* 14) with the implementation available in the BioConductor package Rtreemix [55, 56]. For CBNs as detailed in [33] I used the software from [6]. I used the same default settings for temp (1) and steps (number of nodes^2^) and started the simulated annealing search for the best poset from an initial linear poset as in [6]. For Progression Networks I used the DiProg program (the method we call DiP) from [12] to fit monotone networks (option “MPN”), choosing the best *k* from 1 to 3 (and the results reported here have *ε* = 0.05). Further details about software versions and parameters used for all methods are provided in *Supplementary Material Section* 4.

Other methods have been described in the literature, but I have not been able to use them here. The method in [35] is too slow (analysis of data sets of 200 cases exceeding 4 hours) to be used if we need to do more than 100,000 analysis, as in this paper. The methods in [7, 25] have no software available. Therefore, this paper includes all currently existing approaches for which software is available.

It is important to mention that the methods used differed greatly in speed: the median and mean execution times, over all 172800 analysis performed by each family of methods, were 0.045 and 0.07 seconds for OT, 3.89 and 12.60 seconds for DiP, but 31 and 1127 seconds for CBN. In addition, DiProg (DiP/DiP-A) currently depends on IBM’s CPLEX ILOG library, which not only is not open source but has a severely restrictive license.

### CBN-A, OT-A, DiP-A

In some cases one or more mutations were present in all or almost all of the samples. Even if these are driver mutations on which all other events depend, events with a frequency of 1 are often removed from the graph (e.g., by the OT method) or placed as nodes that descend directly from Root and that have no descendants. To try to minimize this problem, we can augment the data by adding “pseudosamples” that have no mutations in any gene. Adding “pseudosamples” does not amount to knowing anything about the order of events, nor the truth about which genes are drivers or not (and in the Drivers Unknown scenario I always augmented after the filtering step). Data augmentation only requires being able to differentiate between presence and absence of a genetic alteration, mutation, or aberration, which is always assumed in these analyses. In this paper, “CBN-A”, “DiP-A”, and “OT-A”, refer to using CBN, DiP, or OT on data that has been augmented by adding to it another 10% of samples filled with zeroes (0 is the code that denotes no alteration).

### Analysis

We want to address two questions: a) what procedures (choice of Method, Filtering, S.Time, and S.Type) are “best” (for reconstructing the underlying true graph from the data), so that we can choose a course of action when faced with new data; b) what factors have an important effect on performance, even if they are not under user control, so that future research can focus on them. The first question is most straightforwardly addressed by ranking and comparing (via statistical tests of differences) the available options (combinations of Method, Filtering, and Sampling scheme). The second question is best addressed with statistical modeling that focuses on identifying factors with relevant effects. Of course, results from each type of analysis complement each other. The approaches used below reflect these two questions and are based on very different procedures and assumptions. Below I detail the different analyses, after explaining how performance was measured.

#### Metrics: measuring performance

I consider here that the main goal of most studies is the reconstruction of the topology of the graph, which is what captures the order restrictions [12, 15, 34]. There is no single metric that can fully characterize the performance in this task, and therefore I have used four metrics that capture performance along different dimensions. One is a global score of the difference between the inferred graph and the true graph. The other three are measures of classification or diagnostic performance common in medical testing and machine learning [57,58] that focus on the fractions or proportions across specific rows or columns of the confusion matrix (where entries in that matrix are commonly called “true positives”, “false positives”, “false negatives”, and “true negatives”). Thus, the dimensions measured by each of these four metrics relate to concepts already familiar to researchers, and arguably capture the key features of the methods’ behavior. As we will see below, using these four different metrics is also key to understanding some of the major differences between methods.

**Diff** is the sum of the absolute value of the entries in the matrix of the **Difference between the adjacency matrices** of the true (T) and inferred (F) graphs; this is the square of the “usual” Frobenius norm [59] of that matrix difference, and is the same as the “graph edit distance” of [13]. The **Proportion of False Discoveries (PFD)** is defined as 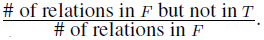.

Following [6, 33], we define “relations” as the transitive closure of “cover relations”. For instance, suppose a graph with *A → B → C* (where *A→ B* means that *A* needs to occur for *B* to occur); the cover relations are *A → B* and *B → C*, but we also include *A → C* in the relation. As in [6, 33], we do not include the root node when finding cover relations and their transitive closure (in contrast to what is done in the computation of Diff). The numerator is, therefore, the number of false positives (FP). The **Proportion of Negative Discoveries (PND)** is defined as 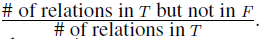 and the numerator is, therefore, the number of false negatives (FN). The **False Positive Fraction (FPF)** is defined as 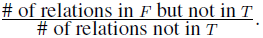 and its numerator is FP. Further details for all metrics are provided in *Supplementary Material Section* 5.

#### Overall ranking of Filtering, Method, and Sampling scheme

To understand what combinations of Method, Filtering, and Sampling scheme are best, I ranked them, averaging the ranks over (subsets of) the other factors (i.e., marginalizing over other factors). As “best” can depend on metric, each metric has to be dealt with separately. More precisely, for each metric separately, I ranked the 36 combinations of Method by S.Time by S.Type (for the Drivers Known scenario) or the 144 combinations of Filtering by Method by S.Time by S.Type (for the Drivers Unknown scenario) in each of the 144 factor combinations defined by True Graph by Model by sh by S.Size. Thus, for each metric, there were 144 separate rankings (where we ranked 36 items in the Drivers Known scenario and 144 items in the Drivers Unknown scenario). Ranking was done using the median (over the 20 replicates) of the metric. Then, for each metric, I obtained the average rank over subsets of the 144 combinations and the averaged ranks were then ranked to obtain the final rankings for each metric (e.g., Table 3 shows, on the left four columns, the ranked average ranks over the 72 combinations of Number of Nodes by Model by sh by S.Size when there are conjunctions and, on the right, the ranks over those 72 combinations when there are no conjunctions). Similar to [60], because the maximum and minimum ranks may differ between scenarios, the ranks were normalized before averaging.

**Table 3.** Overall ranking of all 36 combinations of method and sampling when Drivers are Known with respect to each performance metric. Methods have been ordered by their performance in the first metric. Best 5 methods are shown in bold. Only methods that are within the best 5 in at least one metric are shown (full table as well as tables split by S.Size are available from the Supplementary Material).

| Method and sampling | Conjunction |  |  |  | No conjunction |  |  |  |
| --- | --- | --- | --- | --- | --- | --- | --- | --- |
|  | Diff | PFD | PND | FPF | Diff | PFD | PND | FPF |
| OT-A, last, singleC | **1** | **2** | 14 | 10 | **1** | **3** | **2** | 15 |
| OT-A, last, wholeT_0.5 | **2** | **1** | 15 | 12 | **2** | **4** | **3** | 14 |
| OT-A, last, wholeT_0.01 | **3** | **3** | 7 | 22 | **3** | 6 | **1** | 22 |
| OT-A, unif, singleC | **4** | 6 | 19 | 13 | **4** | 9 | 12.5 | 9.5 |
| OT, unif, singleC | **5** | **5** | 20 | 11 | **5** | 7 | 12.5 | 9.5 |
| OT-A, unif, wholeT_0.01 | 8 | 11 | 16 | 24 | 8 | 11 | **5** | 24 |
| OT, last, singleC | 10 | 9 | 23 | **1** | 10 | **1** | 17 | **2** |
| OT, last, wholeT_0.01 | 11 | **4** | 18 | 18 | 12 | **5** | 14 | 18 |
| OT, last, wholeT_0.5 | 12 | 7 | 24 | **2** | 11 | **2** | 23 | **1** |
| CBN-A, unif, wholeT_0.01 | 13 | 13 | **1** | 26 | 13 | 13 | **4** | 26 |
| CBN-A, unif, singleC | 14 | 16 | **2** | 28 | 15 | 15 | 9 | 29 |
| CBN-A, unif, wholeT_0.5 | 15 | 18 | **3** | 34 | 14 | 16 | 8 | 31 |
| CBN, unif, singleC | 16 | 17 | **5** | 29 | 17 | 19 | 10 | 34 |
| CBN, unif, wholeT_0.01 | 17 | 14 | **4** | 27 | 18 | 14 | 6 | 27 |
| DiP-A, unif, singleC | 31 | 28 | 31 | **4** | 31 | 30 | 31 | 6 |
| DiP, last, wholeT_0.5 | 33 | 35 | 34 | **5** | 34 | 34 | 36 | **5** |
| DiP, unif, singleC | 35 | 31 | 33 | **3** | 35 | 32 | 33 | **3** |
| DiP, unif, wholeT_0.5 | 36 | 36 | 36 | 6 | 36 | 36 | 35 | **4** |

This procedure does not take into account the repeated usage of the same data set for each of the four/sixteen methods, but we are using it simply to rank alternatives. The advantage of this procedure is that it provides an overall view of the results that is equivalent to examining all possible interactions of Filtering by Method by Sampling scheme, marginalizing over all other terms, and this is done with a simple procedure that does not depend on additional modeling assumptions. The disadvantage is that it does not allow us to judge the relative size of different effects.

#### Best subsets of Method and Filtering: within-data set testing

As mentioned above, for each combination of True Graph, Model, sh, S. Size, S. Time and S. Type, I analyzed twenty replicate data sets, and each data set was subject to each Method or each Filtering by Method combination. This design allows us to use paired tests within each among-data set combination to compare each policy (where a policy is a Method in the Drivers Known scenario or a Filtering by Method combination in the Drivers Unknown scenario) with the rest of policies. Separately for each metric and for each of the 864 among-data set combinations, I declared that policy *i* was better (had a smaller metric) than policy *j* if a paired Wilcoxon test had, for the alternative hypothesis of *i < j*, a one-sided *p-value* < 1 − (1 − 0.05)^(1*/numcomps*)^, where *numcomps* is the number of comparisons, or the number of other policies each policy is compared against. This procedure is a simple adaptation (to a paired Wilcoxon test) of the “subset selection” procedures in the analysis of simulation experiments common in industrial settings [61, 62]. For metrics Diff, PND, and FPF, *numcomps* is always five in the Drivers Known case and 23 in the Drivers Unknown case; for PFD sometimes fewer tests were performed if all 20 observations of a policy had missing values (due to not making any discovery). After all pairs of combinations of policies had been compared, for each policy I counted in how many cases it was significantly better than the other policies. The subset of best policies is/are the policy(es) with the largest count. If there is a single policy with that largest number of significant differences, the best subset will be composed of only one policy; otherwise, the best subset will have more than one member (meaning that we cannot tell which of those policies is better using this procedure). The best subsets for each of the 864 among-data set combinations are shown in “best-subsets.pdf”. From these, we can then find the frequency of the different best subsets for selected combinations of factors as shown, for example, in Table 4. A detailed example of this procedure is provided in *Supplementary Material Section* 6.

#### Generalized linear modeling of performance metrics

The procedures above do not provide a direct way to compare the relative magnitude of the effects of different factors. We can approach this problem using a statistical model for each of the metrics. I used generalized linear mixed models (GLMM), where data set was a random effect and the rest of the factors in the design were regarded as fixed effects. Diff was modeled as a Poisson-distributed count, and PND, PFD, and FPF, were modeled as binomial data. Some factors, specially Number of Nodes but possibly also Model, might be regarded as random effects, but I modeled them as fixed effects since the number of different levels is too small (but see *Supplementary Material Section* 7). Data for the Drivers Known and Drivers Unknown scenarios were modeled separately. Models were fitted using INLA [63, 64], a Bayesian approach that uses nested Laplace approximation as an alternative to Markov Chain Monte Carlo (MCMC), with the R package R-INLA. I run all models with two different priors and for model validation I used the cross-validated probability integral transform (PIT) [65] and a simple comparison of fitted vs. observed values. In addition, I also fitted the additive and two-way interaction models for the Drivers Known scenario, and the additive model for the Drivers Unknown scenario, using the R package MCMCglmm [66], with three chains per model and variable; MCMCglmm differs from INLA not only on the use of MCMC (vs. Laplace approximations) but also on the priors and in the inclusion of an observation-level random effect. There were no relevant differences in results.

**Table 4.** Frequencies of best subsets for all metrics when Drivers are Known. The table shows the frequency of the most common best subset combinations. Combinations not shown have a frequency less than 0.05 for all columns. Frequencies normalized by column total (N = 432).

| Best subsets | Conjunction |  |  |  | No conjunction |  |  |  |
| --- | --- | --- | --- | --- | --- | --- | --- | --- |
|  | Diff | PFD | PND | FPF | Diff | PFD | PND | FPF |
| OT, OT-A | 0.63 | 0.49 | 0.05 | 0.06 | 0.66 | 0.55 | 0.27 | 0.08 |
| DiP, DiP-A, OT, OT-A | 0.02 | 0.12 | - | 0.56 | 0.03 | 0.18 | 0.05 | 0.56 |
| CBN, CBN-A | 0.01 | 0.04 | 0.55 | - | - | 0.01 | 0.27 | 0.01 |
| DiP, DiP-A | - | 0.03 | 0.01 | 0.16 | 0.01 | 0.02 | - | 0.14 |
| CBN, CBN-A, OT, OT-A | 0.02 | 0.01 | 0.06 | - | 0.02 | 0.03 | 0.12 | - |
| CBN-A | 0.04 | 0.01 | 0.10 | - | - | 0.01 | 0.03 | 0.01 |
| CBN | - | 0.01 | 0.07 | - | - | - | 0.03 | - |
| OT-A | 0.16 | 0.09 | 0.07 | - | 0.15 | 0.03 | 0.14 | - |
| OT | 0.08 | 0.05 | - | 0.01 | 0.04 | 0.03 | - | - |
| DiP, OT | - | 0.04 | - | 0.08 | 0.01 | 0.06 | - | 0.07 |

Models were fitted using sum-to-zero contrasts: each main effect parameter is to be interpreted as the (marginal) deviation of that level from the overall mean, and the interaction parameter as the deviation of the linear predictor of the cell mean (for that combination of levels) from the addition of the corresponding main effect parameters. As explained in *Supplementary Material Section* 7) we will focus on models with two-way interactions. We will refer to these analyses as the GLMM fits. Further details of the statistical modeling and interpretation of coefficients are provided in *Supplementary Material Section* 7.

## Results

We first examine the results when Drivers are Known. When the identity of drivers is not known (Drivers Unknown), we need to add the step of filtering or selecting mutations before inferring the restrictions.

### Drivers Known

There was wide variation in performance: under some Models and with some Methods perfect results were obtained but, for those same Models and S.Sizes, there were choices of Method and Sampling scheme that led to very large errors, with PFD and PND of 0.7 to 0.9 (see examples of inferred graphs in *Supplementary Material Section* 8 and median values for all metrics for all combinations of factors in *Supplementary Material Section* 16). This emphasizes that even for the easiest models and largest sample sizes, careful choice of Method can be crucial. There was also a large difference in the number of relations inferred (the transitive closure of the cover relations), whether correct or incorrect: the mean values were 19.1, 18.5, 1.7, 2.6, 7.0 and 8.4 for CBN, CBN-A, DiP, DiP-A, OT, and OT-A, respectively (see *Supplementary Material Section* 16).

Table 3 shows the overall ranking of Method and Sampling scheme. OT and OT-A were the best methods according to Diff and PFD. CBN and CBN-A were among the best methods according to PND (in graphs with conjunctions) and DiP and DiP-A according to FPF. This is coherent with the patterns of number of edges (number of dependency relations) inferred: CBN and CBN-A inferred more edges and thus the number of false negatives (FN) decreased, so they had larger sensitivity or recall. But this was done at the cost of increasing the false positives (FP) and, thus, increasing PFD and FPF: a larger fraction of the discoveries were false (precision was smaller) and a larger fraction of the non-existing relationships were regarded as being present (specificity was smaller). DiP and DiP-A showed the opposite trend: these were the methods that inferred the smallest number of relations (in many cases no edges, beyond those from Root, were inferred), leading to a smaller number of false positives (FP), so that a smaller fraction of non-existing relations were regarded as being present, but this was done at the cost of a very large number of FN that affected not only PND but also Diff.

Figures 2 and 3 show the coefficients from the GLMM fits. From Figure 2 we see that DiP and DiP-A only performed well with respect to metric FPF, and CBN and CBN-A only with respect to metric PND. However, for metric PND the better performance of CBN/CBN-A compared to other methods was concentrated in graphs with conjunctions. The left column for PND in Table 3 shows that the best five methods were all CBN/CBN-A, but the right column for PND shows that OT-A occupies the first three positions. The analysis of frequencies of best subsets, in Table 4, again reveals the same patterns: OT and OT-A were clearly the best methods for metrics Diff and PFD, and were best methods with DiP/DiP-A for metric FPF. CBN and CBN-A were in best subsets that did not include any of the other methods in 55% of the cases for metric PND in graphs with conjunction. In the absence of conjunctions, however, the frequency of best subsets of CBN/CBN-A was the same as that of OT/OT-A, and subsets that did not include CBN/CBN-A were more prevalent than those that included CBN/CBN-A (and this association between subset and conjunction was highly significant, *p <* 2.2 * 10*^-^*^16^, from a chi-square test). That the best performance of OT/OT-A in graphs with conjunctions cannot be perfect should be expected, and we should only see perfect performance in these cases, if at all, with CBN/CBN-A or DiP/DiP-A. Two extreme cases (which also provide an internal consistency check) are graphs “7-A” and “11-A” (both have conjunctions): perfect performance was achieved for the first with CBN-A and for the second with DiP (S.Size = 1000, McF_6, sh Inf and 0, S.Time unif and last, respectively —see *Supplementary Material Section* 16).

A simple marginal plot is shown in Figure 4 which provides a view of the above results in a scale that is directly interpretable, and which also illustrates the interaction Method by Conjunction that we see in Figure 2. Conjunctions degraded performance for Diff for all methods, and were irrelevant for FPF; for the other metrics their effect was method-dependent. OT and OT-A were better for metrics Diff and PFD regardless of the presence/absence of conjunctions, and were essentially as good as DiP/DiP-A for FPF. For PND, when there were conjunctions, CBN/CBN-A were the best methods, but when there were no conjunctions OT-A was the best. Interestingly, for CBN/CBN-A, PFD was smaller in the presence of conjunctions (an effect that we can also see in Figure 2), probably due to the tendency of CBN/CBN-A to infer an excess of conjunctions (see figures of reconstructed trees in *Supplementary Material Section* 8).

**Figure 3.**
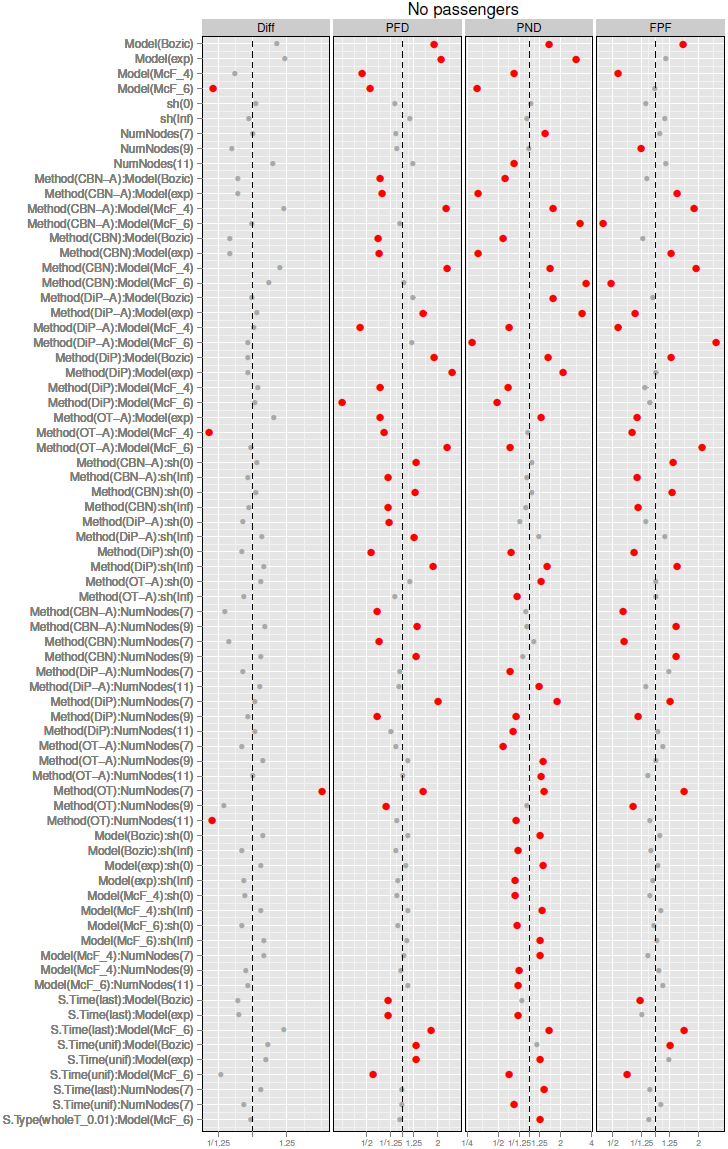
Drivers Known, plot of the coefficients Model, sh, Graph, and their interactions with all other terms from the GLMMs for each of the metrics. See legend for Figure 2.

**Figure 4.**
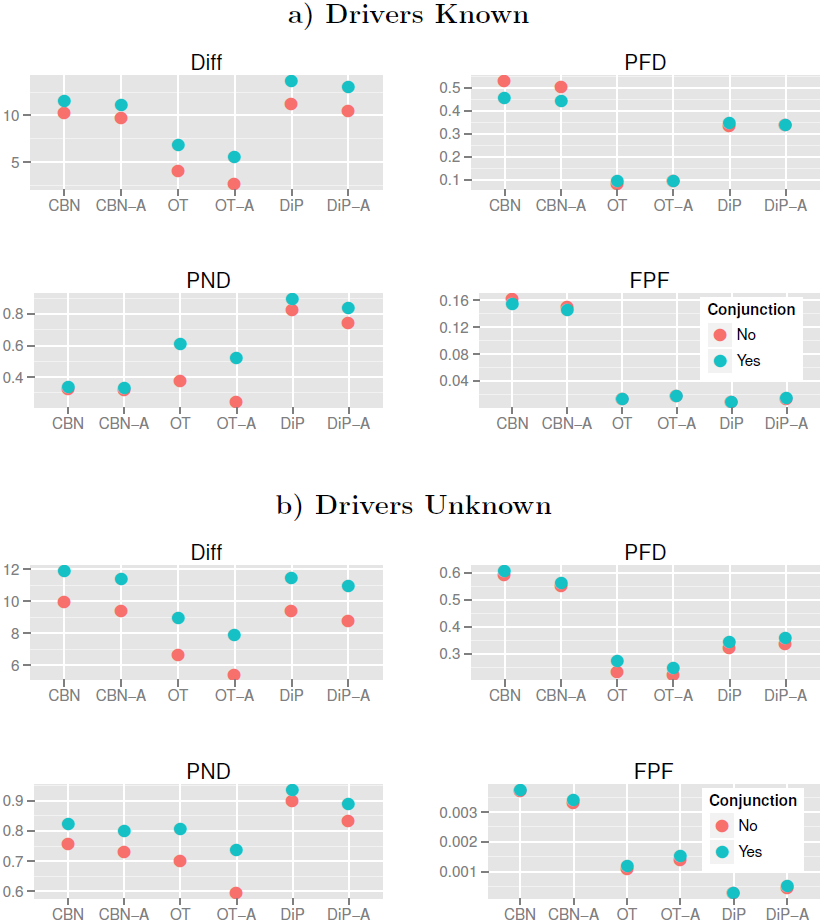
Mean number of each of the metrics for the different combinations of Method and Conjunction in (a) the Drivers Known and (b) the Drivers Unknown scenarios. Y-axis is in the scale of the variable (fractions for PFD, PND, FPF and sum of differences for Diff). Each mean value shown is the mean of 8640 and 34560 values for the Drivers Known and Unknown, respectively.

Figure 2, as well as Table 4 and Table 3 (full tables available in Supplementary Material) show that OT-A was generally superior to OT, and similar but weaker patterns affected CBN and DiP. The magnitude of the differences between augmented and non-augmented alternatives, however, depended strongly on metric and showed interactions with other effects (e.g., S.Time). If we focus on Diff, with OT and DiP there was little to loose from always using the augmented alternative, even if no mutation had a frequency of one, but this was not the case with CBN (see *Supplementary Material Section* 13).

The effect of S.Time was marginally very small (Figure 2), and there were also small interactions with Method, which depended on metric (see *Supplementary Material Section* 9). These differences in performance of each Method with different S.Time in each metric can also be observed in the overall ranking of methods (Table 3), as well as in the change in the relative frequencies of OT, OT-A and CBN in best subsets as a function of S.Time (see *Supplementary Material Section* 11). All of these interactions, however, were much smaller than the main effect of Method and should rarely affect which method to choose.

Although the marginal effects of S.Type were generally small (and single cell sampling provided no benefit), overall it seems best to avoid whole tumor sampling with very small detection thresholds. However, as with many other effects, this was reverted with PND. This result is intuitively reasonable: whole tumor sampling at very low thresholds can lead to obtaining samples where we observe together two low frequency events that rarely occur together in the same individual clone (i.e., that do not correspond to a pattern encoded in the true graph), leading to the observation of possible artifacts, but also allowing the detection of co-occurring events of very low frequency. The effects of S.Type were amplified by method (the interaction of Method by S.Type in Figure 2), and this effect was strongly metric-dependent (see also *Supplementary Material Section* 9). The marginal effect of S.Size was as expected: larger sample size led to better performance with all metrics. However, the effect in performance was small compared to the effects of choosing a bad method or even the effects of S.Type and S.Time for some of the metrics (or the effects of non user-controllable factors such as Model). Moreover, the effects of increases in sample size depended on method and metric (see also *Supplementary Material Section* 9): DiP/DiP-A were, comparatively, the methods that benefited the most from increasing S.Size (except for FPF).

Regarding variables that are not under user control, Model and its interactions with other factors had a strong effect on performance (see Figure 3). Overall, the McFarland models led to better performance (see also Figure 5). The differences between evolutionary models also explained the interaction S.Time by Model. S.Time = uniform benefited especially McF_6 (whereas the opposite trend was observed with Bozic and exp; see also *Supplementary Material Section* 9). Due to the strong density dependence of McF_6, if we sample at the end it will not be easy to observe intermediate steps that involve only a few mutations, since the final population will be composed of clones with five to six driver mutations.

**Figure 5.**
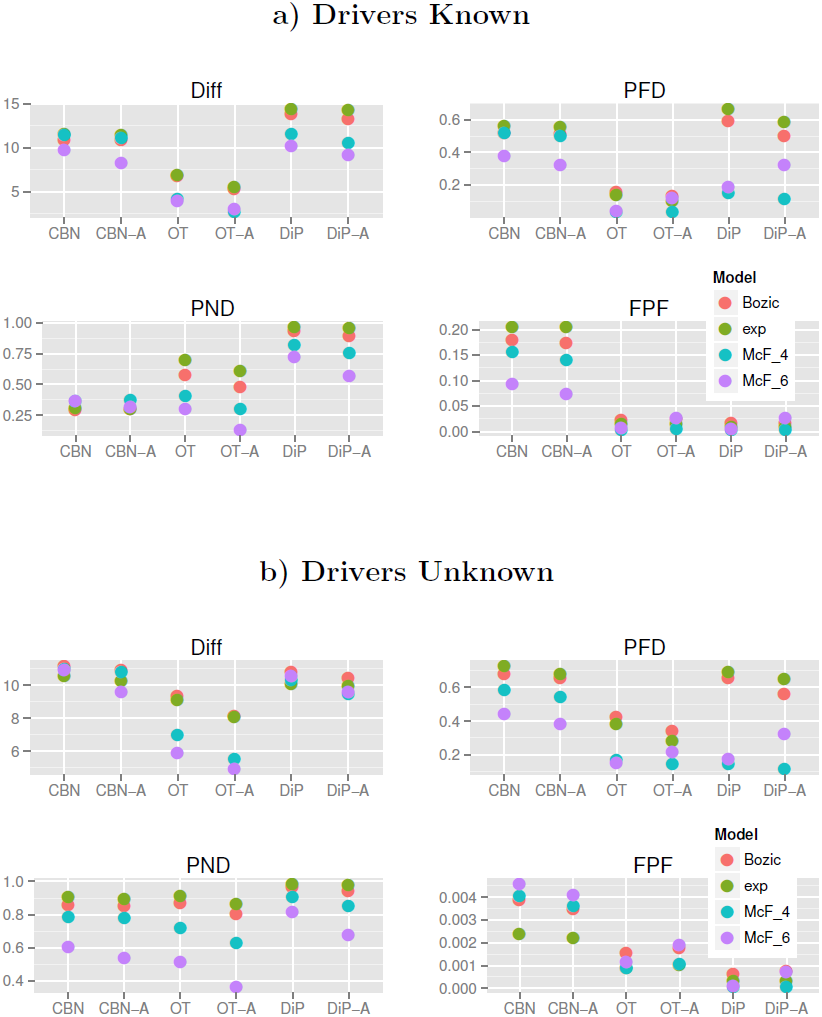
Mean of each of the metrics for the different combinations of Method and Model in (a) the Drivers Known and (b) the Drivers Unknown scenarios. Each mean value shown is the mean of 4320 and 17280 values for the Drivers Known and Unknown, respectively.

Model also showed interactions with Method, and Model determined the effects of sh (the strength of enforcement of monotonicity) and affected its interaction with Method. If we knew that the true model was McF_6, even when minimizing PND we should then choose OT-A (or even OT), not CBN or CBN-A; this can be seen from the marginal results displayed in Figure 5 (or from the coefficients in Figure 3, or by adding the relevant coefficients from the model fit of PND in *Supplementary Material Section* 12). Conversely, for PND, the performance of CBN/CBN-A is much more robust to evolutionary model variation.

Method and sh also showed interactions (Figure 6) and the performance of DiP and DiP-A improved with sh = 0, which contrasts with CBN/CBN-A and OT/OT-A (where sh = Inf leads to better performance): all three families of procedures make allowance for deviations from monotonicity, but the model behind DiP was able to deal with (or even be favored by) them better. Finally, regarding the interaction Model by sh, from Figure 3 and Figure 7 we see that with Bozic and exp, sh = 0 consistently led to worse performance over the four metrics, but it had the opposite effect on the McF models. This is understandable, since the McF models have very strong density dependence of fitness: if the graph specifies *A → B* and a clone has B without A, even if there is no explicit penalty via the birth rate (i.e., sh = 0), we will be unlikely to observe it, since it will be under a severe relative disadvantage compared to clones with A and not B, and under a much more severe disadvantage compared to clones with both A and B. Thus, even if sh = 0, the McF models by their very nature intrinsically incorporate a strong penalty for any mutation order that does not strictly conform to that encoded in the true graph. In other words, genes that can act as drivers or passengers depending on genetic context are much less likely to be observed in their passenger role in the McF models. As we can see, therefore, differences in evolutionary model can modify how deviations from monotonicity affect the performance of different methods and these results underline the importance of explicitly considering evolutionary model and deviations from monotonicity.

**Figure 6.**
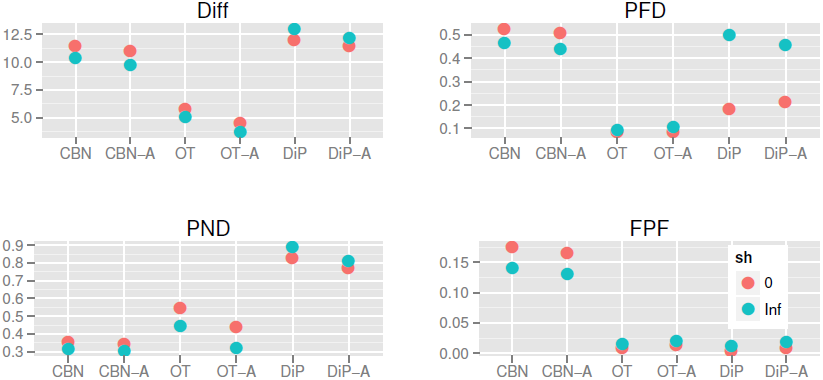
Mean number of each of the metrics in the Drivers Known scenario, for the different combinations of Method and sh. Each value shown is the mean of 8640 values.

**Figure 7.**
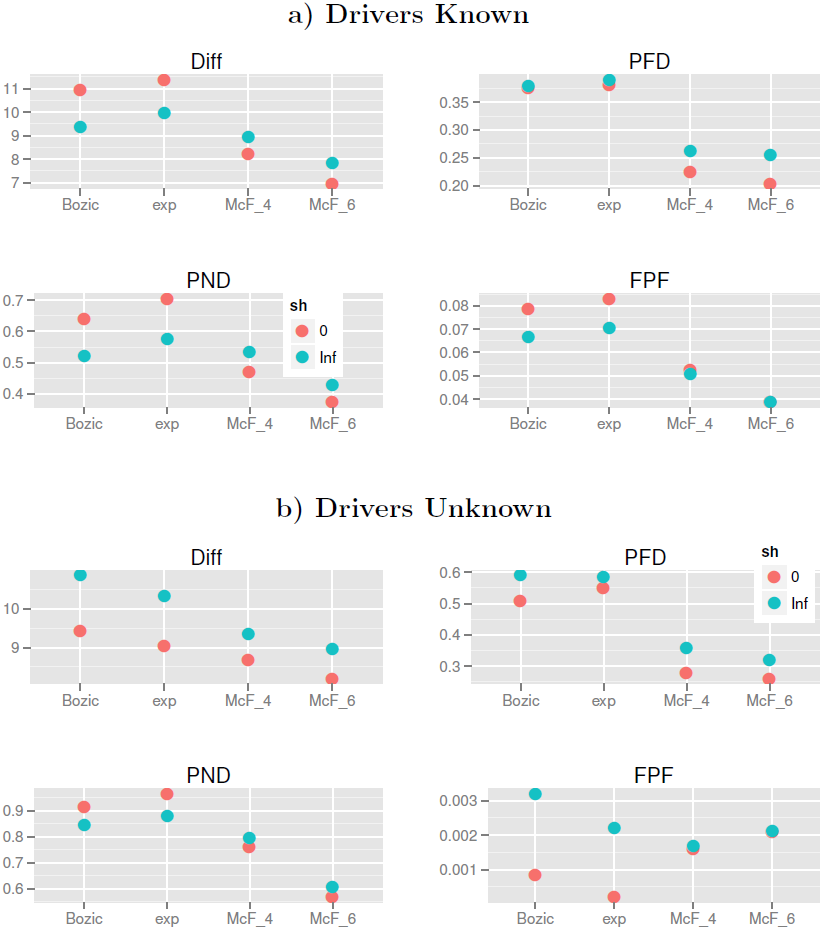
Mean number of each of the metrics for the different combinations of Model and sh in (a) the Drivers Known and (b) the Drivers Unknown scenarios. Each value shown is the mean of 12960 and 51840 values for the Drivers Known and Unknown, respectively.

Finally, Graph had an effect, which for some metrics was large, and also presented large interactions with Model, Method and S.Time for some metrics; these effects will not be examined here further (although a specific one will be addressed in the Discussion), but it is important to notice that the graphs used in this study differed in many ways that are not simply summarized by the number of nodes (see *Supplementary Material Section* 3).

### Drivers Unknown

Many results here were similar to the ones in the previous section, so we will focus on the main differences as well as the added factor of filtering. Overall, performance was worse when Drivers were Unknown (the overall mean values for Diff, PFD, PND, and FPF were 9.2, 0.30 0.53, 0.06 when Drivers were Known, but 9.4, 0.41, 0.80, and 0.002 when Drivers were Unknown —FPF is not comparable because of the way TN has to be calculated when Drivers are Unknown; see *Supplementary Material Section* 5). As those number show, the main problem was the large increase in PND, or failing to detect existing edges: see panels a) and b) of Figure 4 for a graphical comparison. We can see in Figure 8 that filtering almost always resulted in selecting a number of genes smaller than the true number of nodes in the true graph. However, there were cases when performance was perfect or almost perfect for all metrics (trees without conjunctions, with model McF_6, method OT-A, Dip-A, and occasionally OT, and filtering S5 and rarely S1 or J1; see *Supplementary Material Section* 16.2).

**Figure 8.**
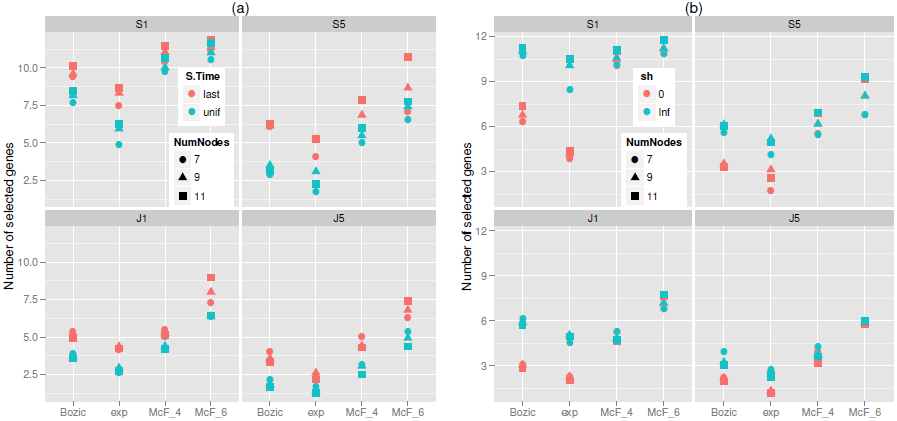
Mean number of genes selected for the different combinations of Model and Filter by S.Time (panel (a)) or sh (panel (b)). Different symbol shapes identify the number of true nodes (NumNodes) of the true graph. Note that the number of genes selected is a function of Filtering, not Method. Each value shown is the mean of 720 values.

Figures 9 and 10 show the coefficients from the GLMM fits. The marginal effect of Filtering (Figure 9) was as expected: more stringent filtering (J5) decreased FP and thus led to better performance for both PFD and FPF, but less stringent filtering (S1) was better for missing fewer patterns, and thus led to better performance in PND (where J5 shows terrible performance); this pattern is also seen in Figure 11. A reasonable overall choice is probably S5: it was the best filtering for the Diff metric, and did reasonably well for all the other metrics. However, as we have already seen repeatedly, the choice of the best filtering is metric-dependent, as we can also see from the overall ranking of methods (Table 5).

**Figure 9.**
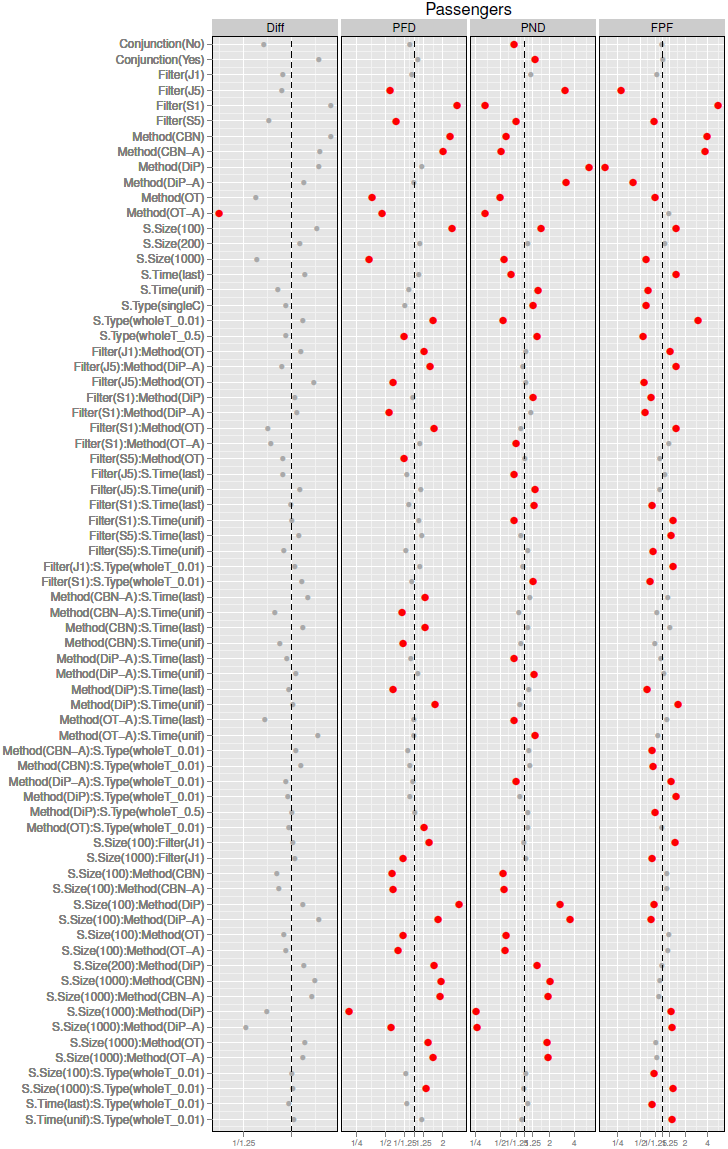
Drivers Unknown, plot of the coefficients for Conjunction, Filtering, Method, S.Time, S.Type and S.Size from the GLMMs for each of the metrics. See legend for Figure 2.

**Figure 10.**
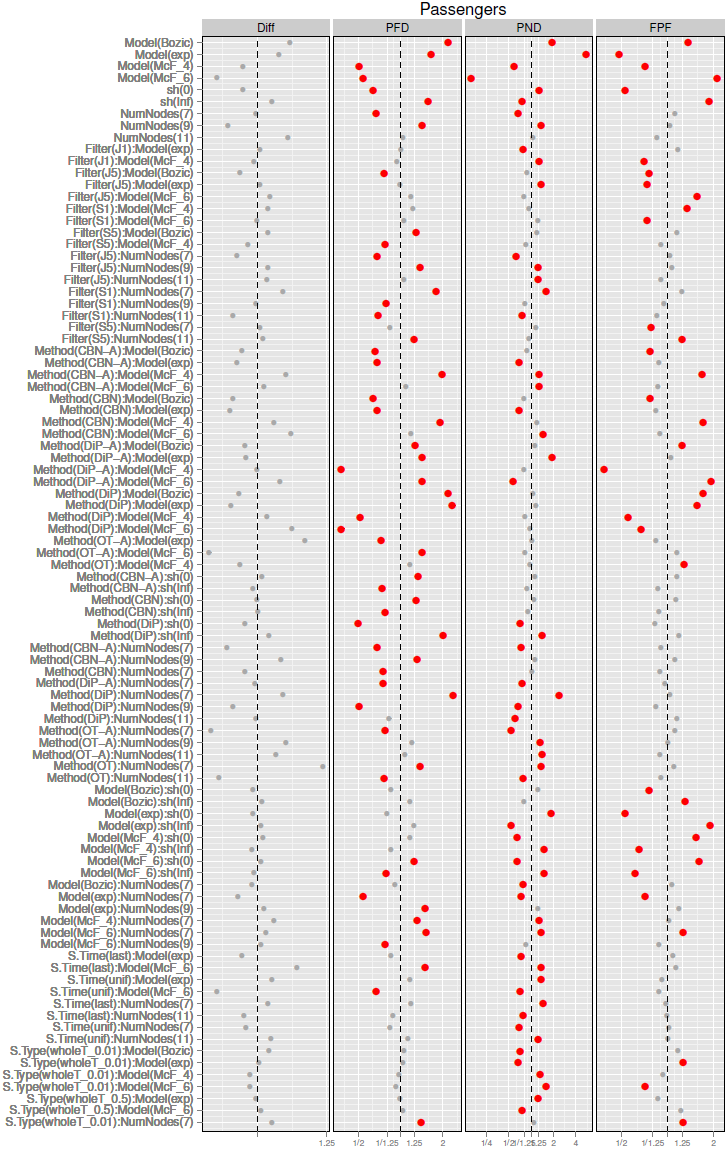
Drivers Unknown, plot of the coefficients Model, sh, Graph, and their interactions with all other terms from the GLMMs for each of the metrics. See legend for Figure 2.

**Figure 11.**
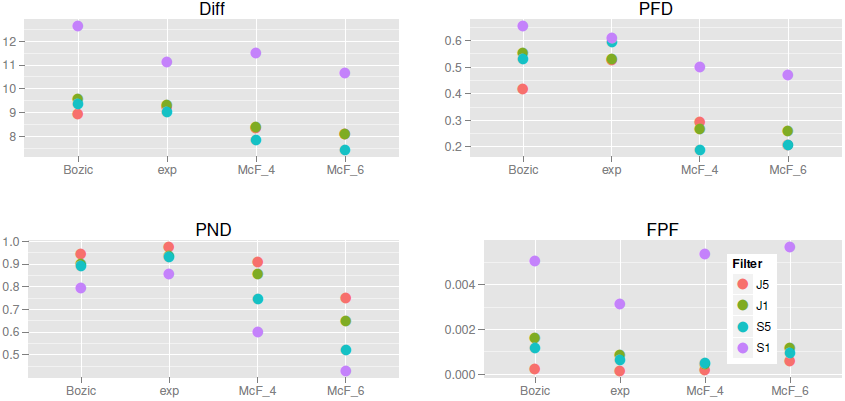
Mean of each of the metrics in the Drivers Unknown scenario showing the interaction of Model by Filter. Each value shown is the mean of 25920 values.

**Table 5.** Overall ranking of all 144 combinations of method, filtering, and sampling when Drivers are Unknown with respect to each performance metric. Methods have been ordered by their performance in the first metric. Best 10 methods are shown in bold. Only methods that are within the best 10 in at least one metric are shown (full table as well as tables split by S.Size are available from the Supplementary Material).

| Method and sampling | Conjunction |  |  |  | No conjunction |  |  |  |
| --- | --- | --- | --- | --- | --- | --- | --- | --- |
|  | Diff | PFD | PND | FPF | Diff | PFD | PND | FPF |
| S5, OT-A, last, singleC | **1** | **7** | 15 | 60 | **1** | 13 | 13 | 60 |
| S5, OT-A, last, wholeT_0.5 | **2** | **6** | 23 | 59 | **2** | **7** | 23 | 59 |
| J5, OT-A, last, wholeT_0.01 | **3** | **1** | 26 | 71 | **4** | **2** | 29 | 74 |
| S5, OT-A, last, wholeT_0.01 | **4** | **2** | **2** | 94 | **3** | **5** | **4** | 94 |
| S5, OT-A, unif, wholeT_0.01 | **5** | 17 | 33 | 69 | **10** | 17 | 32.5 | 68.5 |
| S5, OT, unif, singleC | **6** | 41 | 61.5 | 28 | **9** | 38 | 55.5 | 20.5 |
| S5, OT-A, unif, singleC | **7** | 50 | 61.5 | 31 | **8** | 49 | 55.5 | 20.5 |
| J1, OT-A, last, singleC | **8** | 15 | 38 | 80 | **6** | 12 | 37 | 75 |
| J1, OT-A, last, wholeT_0.5 | **9** | **10** | 39 | 79 | **7** | **6** | 39 | 71 |
| S5, OT-A, unif, wholeT_0.5 | **10** | 47 | 67 | 29.5 | 14 | 45 | 60.5 | 27.5 |
| S5, OT, unif, wholeT_0.01 | 11 | 11 | 34 | 68 | 11 | **10** | 32.5 | 68.5 |
| J1, OT-A, last, wholeT_0.01 | 13 | 21 | **9** | 109 | 15 | 19 | **8** | 108 |
| S1, OT-A, last, wholeT_0.5 | 18 | 39 | **3** | 117 | **5** | 27 | **3** | 116 |
| S1, OT-A, last, singleC | 21 | 38 | **4** | 120 | 12 | 32 | **2** | 120 |
| S5, OT, last, singleC | 23 | 16 | 51 | 21 | 20 | **8** | 42 | 19 |
| S5, OT, last, wholeT_0.5 | 24 | **3** | 55 | 16 | 23 | **3** | 49 | 17 |
| J1, OT, unif, wholeT_0.01 | 29 | **9** | 36 | 91 | 30 | 14 | 36 | 91 |
| S5, CBN-A, unif, wholeT_0.01 | 30 | **4** | 37 | 89 | 40 | 20 | 40 | 103 |
| J5, OT, last, wholeT_0.01 | 31 | **5** | 59 | 32 | 36 | **4** | 62 | 50 |
| S1, OT-A, last, wholeT_0.01 | 36 | 48 | **1** | 134 | 24 | 50 | **1** | 134 |
| S5, OT, last, wholeT_0.01 | 38 | **8** | 29 | 83 | 34 | **1** | 24 | 82 |
| J1, OT, last, wholeT_0.5 | 47 | 14 | 89 | 61 | 38 | **9** | 83 | 43 |
| J5, OT, last, singleC | 48 | 35 | 101 | **4** | 58 | 43 | 109 | **4.5** |
| J5, DiP-A, unif, singleC | 49 | 125 | 139 | 10.5 | 51 | 128 | 138 | **9** |
| J5, OT, last, wholeT_0.5 | 57 | 20 | 102 | **4** | 75 | 23 | 112 | **4.5** |
| J5, DiP, unif, singleC | 61 | 131 | 143 | **8** | 60 | 132 | 144 | **4.5** |
| J5, DiP, unif, wholeT_0.5 | 64 | 144 | 144 | **4** | 64 | 144 | 143 | **4.5** |
| J5, DiP, last, singleC | 79 | 115 | 140 | **4** | 81 | 119 | 140 | **4.5** |
| J5, DiP, last, wholeT_0.5 | 82 | 137 | 138 | **4** | 83 | 136 | 139 | **4.5** |
| S5, DiP, last, wholeT_0.5 | 83 | 134 | 120 | **9** | 87 | 130 | 110 | **10** |
| J1, DiP, last, wholeT_0.5 | 91 | 132 | 134 | **4** | 89 | 134 | 134 | **4.5** |
| J1, DiP, last, singleC | 92 | 110 | 135 | **4** | 93 | 115 | 132 | **4.5** |
| S1, OT-A, unif, wholeT_0.01 | 102 | 65 | **5** | 136 | 70 | 59 | **5** | 136 |
| S1, OT, unif, wholeT_0.01 | 109 | 62 | **6** | 135 | 74 | 55 | **6** | 135 |
| S1, CBN-A, unif, wholeT_0.01 | 137 | 82 | **7** | 143 | 140 | 81 | **7** | 143 |
| S1, CBN-A, last, wholeT_0.01 | 142 | 93 | **10** | 137 | 139 | 94 | 12 | 137 |
| S1, CBN, unif, wholeT_0.01 | 143 | 84 | **8** | 144 | 143 | 90 | **9** | 144 |

One difference with the results when Drivers are Known was related to Method: here, OT/OT-A were almost always better than CBN and CBN-A. This can be seen in Figure 9 —OT and OT-A had much smaller coefficients for all metrics, except for OT under PND—, and comparing panels a) and b) of Figure 4. The same pattern was seen in the overall ranking of methods (Table 5): CBN/CBN-A were rarely among the best performers, not even with conjunctions and with metric PND. Similar results were observed in the analysis of best subsets (Table 6): CBN/CBN-A were rarely among best subsets, and when they were it was generally in best-subsets that included also OT and OT-A, from which they could not be differentiated by the Wilcoxon tests. The results for DiP and DiP-A were similar to the Drivers Known scenario: these methods led to the smallest FPF, but did so at the cost of the other metrics. However, in this scenario DiP and DiP-A were better performers than CBN and CBN-A for Diff.

**Table 6.** Frequencies of best subsets for all metrics when Drivers are Unknown. The table shows the frequency of the most common best subset combinations. Combinations not shown have a frequency less than 0.05 for all columns or are composed of more than 10 individual best methods. Frequencies normalized by column total (N = 432). ‘A:B’ denotes filtering with A and using method B.

| Best subsets | Conjunction |  |  |  | No conjunction |  |  |  |
| --- | --- | --- | --- | --- | --- | --- | --- | --- |
|  | Diff | PFD | PND | FPF | Diff | PFD | PND | FPF |
| S1:OT-A, S5:OT-A | 0.01 | - | 0.07 | - | 0.01 | - | 0.08 | - |
| S1:OT, S1:OT-A | 0.10 | 0.10 | 0.23 | - | 0.13 | 0.11 | 0.35 | - |
| S1:OT-A | 0.02 | - | 0.06 | - | 0.02 | - | 0.06 | - |
| S5:OT-A | 0.07 | - | - | - | 0.09 | - | - | - |
| S5:OT | 0.06 | - | - | - | 0.05 | - | - | - |
| S5:OT, S5:OT-A | 0.20 | 0.05 | - | - | 0.22 | 0.07 | - | - |
| S5:DiP, S5:DiP-A, S5:OT, S5:OT-A | 0.06 | - | - | - | 0.07 | - | - | - |
| S1:OT, S1:OT-A, S5:OT, S5:OT-A | 0.01 | - | 0.03 | - | 0.03 | - | 0.07 | - |
| J5:DiP, J5:DiP-A, J5:OT, J5:OT-A | - | 0.01 | - | 0.06 | - | - | - | 0.05 |
| J5:OT, J5:OT-A, S5:OT, S5:OT-A | - | 0.06 | - | - | 0.01 | 0.04 | - | - |
| S1:CBN, S1:CBN-A, S1:OT, S1:OT-A | - | - | 0.18 | - | - | - | 0.12 | - |

There were interactions of Filter by Method (see figures in *Supplementary Material Section* 9), and their direction and magnitude depended on metric, but these interactions were not large enough to revert our preferences of methods. S.Type followed similar patterns as seen for the Drivers Known scenario, although the effects were of larger magnitude, and its interactions with Method or Filter (see also *Supplementary Material Section* 9), even when present, produced no change in the ordering of preferences of Filtering, Method, or S.Type. The effects of S.Size or its interactions with Method were similar to those in the Drivers Known scenario, although S.Size seemed to be more important for decreasing PFD when Drivers are Unknown (figures in *Supplementary Material Section* 9).

Other effects and interactions were different with respect to the Drivers Known scenario, illustrating that the need to filter passengers can lead to counterintuitive and much more complicated interpretation of results. There was an interaction of Filter by Model (Figure 10): the best choice of S5 vs. J5 for metrics Diff and PFD depended on the Model, as shown in Figure 11. Moreover, we can see that the very poor results in metric FPF of model McF_6 were largely due to its terrible results with Filter S1 (see also below). There was also an interaction Model by Method, and OT/OT-A were now superior in all metrics (except FPF) not only overall, but also virtually for every one of the four models, as we can see from panel b) in Figure 5.

The effect of S.Time for all metrics except Diff was reverted and had larger magnitude compared to the Drivers Known scenario. The cause is that S.Time affected the number of genes that were selected and thus the number of false positives (FP) and false negatives (FN). As seen in panel (a) of Figure 8, under S.Time = uniform fewer genes were selected for all filtering methods (the number of genes selected is independent of Method but depends on Filtering). Under uniform sampling many of the samples have few mutations (those corresponding to the early stages of the disease), and thus fewer genes are above the filtering thresholds. As fewer genes are selected under S.Time = uniform, the number of FP goes down, and thus both PFD and FPF showed a decrease (and the opposite pattern was observed for PND). None of these phenomena were present when Drivers are Known, because there was no filtering step there. For Diff, however, the marginal effect of S.Time = uniform was still slightly better than that of last-period sampling, as was when Drivers are Known. These differences of the effect of S.Time between the Drivers Known and Drivers Unknown scenarios explain also the differences in the patterns of interactions of Method by S.Time and Model by S.Time (see figures in *Supplementary Material Section* 9).

Two other differences with the Drivers Known scenario are the poor performance of model McF_6 with metric FPF, and some of the interactions between Model and sh in several metrics, and both probably are related by the filtering step. The poor performance of McF_6 in FPF was not paralleled by a poor performance in PFD, which also has FP in the numerator. Thus, per discovery made (i.e., for each relationship found by the methods), McF_6 was actually the model that allowed achieving best results (the best PFD). But the number of errors (FP) increased disproportionately with respect to the number of edges that were not present (the denominator of FPF, a constant for each True Graph). Figure 8 (both panels) shows that, for all filtering methods, McF_6 was the model that resulted in the largest number of genes selected: the strong density dependence of McF_6, specially when S.Time = last, will tend to return samples where most driver mutations are present in many samples. This increase in the number of genes selected can lead to inferring a larger number of relationships and, ultimately, to an increase in FP. This would also explain that sh = 0, specially with PFD and FPF, led to a large decrease in error rates. Panel (b) of Figure 8 shows that, for all filtering procedures, sh = 0 led to smaller number of genes being selected with models Bozic and Exp (but had no effect on the McF models), and Bozic and exp were the two models where sh = 0 led to a larger relative improvement in performance in PFD and FPF (Figure 10 and panel b of Figure 7). Interestingly, the difference between the two levels of sh is more pronounced in S1, and suggests that sh = Inf allows the accumulation of larger numbers of mutated genes that pass the less stringent filters, but that have too little signal for the inference of the restrictions (in contrast, sh = 0 leads to the accumulation of mutations with too low a frequency to even pass the filters). That sh should have virtually no effect on number of genes in the McF models has been explained before. These interactions, therefore, highlight the complex and counterintuitive relationships between Model, sh, and Filtering, that then cascade onto the overall performance differences between Methods or Sampling schemes.

## Discussion

This paper presents a comprehensive study that has examined, for the first time, the effects of sampling decisions, evolutionary models, and presence of passenger mutations in the performance of methods for the inference of restrictions in the order of accumulation of mutations.

From the point of view of a user of these methods the main results can be summarized as follows:

1. Method and sampling choice should be guided by the metric considered most important: no combination of Method, Filtering, and Sampling scheme excels in all metrics. This is not unexpected, but it is worth emphasizing that each metric is sensitive to a different kind of deviation, and the results in this paper show that characterizing behavior with only one or two metrics could have been deceptive. Moreover, as we have seen, the relative strengths of each method are better captured and understood using different metrics.
2. In terms of method choice, a very simple summary is (see also examples of inferred graphs in *Supplementary Material Section* 8): CBN tends to return graphs with too many edges, including too many conjunctions, DiP tends to return graphs with too few edges between non-root nodes, and OT does a good overall job even though it will fail, by construction, to return any conjunction. In more detail, OT and OT-A are the best methods, except if we are particularly interested in minimizing PND and we suspect conjunctions are present (when we might want to consider CBN) or FPF (when we might want to consider DiProg). Since it is impossible for OT to return any conjunctions, further research on computationally efficient methods to recover conjunctions is sorely needed.
3. Using frequency-based statistics when we do not know which mutations are true passengers can lead to a heavy performance penalty in the form of failure to discover existing restrictions. In addition, having to filter genes makes it much harder to intuitively understand, and reason about, what is likely to happen in any scenario, and this in turns makes interpretation of results and reconciliation of output from different methods much harder. Thus, it probably pays off to try to use other approaches that incorporate information about non-silent mutation rates, pathway information together with combinatorial properties of drivers in pathwayws, or functional consequences of mutations to differentiate drivers from passengers [67–72]. It might not always be possible to use these other methods. If we need to rely on frequency-based approaches, selecting those mutations with a frequency larger than 5% is an overall reasonable choice (but not the single best choice for any metric other than Diff).
4. Sampling time and type, by themselves, had minor effects compared to, say, filtering or method choice (and we will rarely have control over these factors when we use data already available in databases). However, we might have information about characteristics of the tumor that indicate that it is in an exponential (the Bozic or exp models) or logistic-like growth (the McF models) phase; sampling as late as possible is to be preferred for the first cases, whereas trying to obtain samples distributed over different stages of the disease is best in the second. The best choice of sampling time, however, will depend on metric and whether or not we are certain about which are the driver genes.
5. Single cell sampling is about as good as whole tumor sampling, unless we use whole tumor sampling with extremely small detection thresholds (which leads to poorer performance, except for metric PND).
6. Although with S.Size the larger the better, its effect is relatively minor for OT and CBN (not so for DiP), a result that agrees with those in [13], specially for some metrics. In particular, resources might be better spent trying to be certain about the identity of the true drivers than increasing sample size from 100 to 1000.
7. Data augmentation is not always the right choice, although with OT and DiP there is little to loose from always using data augmentation but potentially a lot to loose from not using augmentation (see *Supplementary Material Section* 13). Unfortunately, simple rules of thumb like “always use data augmentation when at least one gene has a frequency of one, and never otherwise” do not work well, especially across all methods, and appropriate choice warrants further study.

Most of the above results cannot be compared to any other studies, since these factors have not been considered before. It has been previously discussed [15, 34] that very good reconstructions of oncogenetic trees are achievable with realistic sample sizes, and we have seen that, at least under several scenarios, it is possible to obtain perfect reconstructions even with sample sizes as small as 100.

The results concerning the superiority of OT with respect to CBN contrast with those of Hainke et al. [13], who find that CBN outperforms OT. A more detailed discussion is provided in *Supplementary Material Section* 14, but these differences are attributable to [13] basing their conclusion on a single graph with a small number of nodes for each scenario. *Supplementary Material Section* 14 shows the within-data set difference in the Diff metric between CBN and OT separately for the different combinations of Model, sh, True Graph, and Sampling scheme (under the Drivers Known scenario only, since [13] do not consider passengers): OT systematically outperformed CBN except for Graph 7A when sampling last (under all models except McF_6), and Graph 7B when sampling last under the Bozic and exponential models. This pattern was reproduced when I fitted the OT models with the Rtreemix package [55] (see *Supplementary Material Section* 14). The single tree used by [13] for the non-conjunction case (their Figure A1) contains five nodes, and the single graph they used for the case with conjunctions contains four nodes (their Figure A7), a very small number of nodes compared to the trees that are seen on the literature (see Material and Methods section). Our graphs 7A and 7B, those where CBN outperformed OT in certain scenarios, are the closest in number of nodes to the graphs in [13].

Given that the results of Hainke et al. [13] are, thus, probably not really contradictory with ours, is the recommendation that practitioners generally use OT instead of CBN still valid? Yes, since most graphs in the literature (including studies involving citogenetic bands and genes) contain many more than four or five nodes, and we cannot be sure if the evolutionary model is one that would favor using CBN. In addition, we have focused only on the comparison between CBN and OT, since those are the only variants used in [13]: if we included OT-A in the comparison, CBN (or CBN-A) would then very rarely be better alternatives (see, e.g., *Supplementary Material Section* 14). However, this apparent difference in results emphasizes the need for considering at least a few different scenarios with regards to potentially key variables, and suggests that a through examination of the impact of graph characteristics (and its interaction with evolutionary model and sampling scheme) on method performance is warranted.

This paper is also one of the first to explicitly connect evolutionary models with restrictions on the order of mutations. Recently, in [41] a simulation tool has been described where restrictions are incorporated into the evolutionary model of [43]; our approach is more general as [41] are limited to four drivers and no passengers whereas, in addition to passengers and other evolutionary models, we can specify restrictions in the order of mutations using arbitrary graphs and allowing for a range of deviations from monotonicity. In fact, one of the attractive features of OTs, CBNs, and Progression Networks is their mechanistic interpretation as graphs that encode restrictions in the order in which driver mutations can accumulate [12, 13, 15, 34]. And one major result of this paper is that inferring those restrictions can be strongly affected by evolutionary model (including deviations from monotonicity) and sampling scheme, and that the relative effect of these factors depends on the metric used. Yet restrictions in the order of driver mutations and evolutionary models are virtually always examined separately.

There is a rich literature about tumor progression models that focuses on the consequences of drivers, passengers, and variation in selection pressures [20,43,44,46,73,74], and a largely separate body of work [13–15, 19, 21, 33, 34, 47] that deals with understanding the restrictions and order of accumulation of mutations (but see [6] for a connection between the *λ_i_* of CBNs and selection coefficients, in the context of the Fisher-Wright model of tumor progression in [20]). The work of Cheng et al. [7, 8] tries to infer the order of mutations within a explicit evolutionary model of tumor progression; unfortunately, no software is available, and thus comparisons are not possible.

Focusing on methods with available software, the actual values (and, thus, interpretation) of the conditional probabilities inferred by OTs or the *λ_i_* parameter for the waiting time to event *i*, in CBNs, will be a complex interplay between the restrictions encoded in the graphs and the details of the tumor progression model as well as the sampling scheme used. For both OTs and CBNs we should expect estimated *λ*s and conditional probabilities to vary by node level or depth (where level or depth refers here to how many edges there are in the path to the root): deeper nodes will show smaller values and, for a given depth, *λ*s/conditional probabilities should be larger for those nodes than “unlock” more downwards mutations. The strength of this effect will increase with the number of nodes along the largest path along the graph, especially when the evolutionary model and sampling scheme result in strong selection for clones with many drivers, and we should see competition between multiple nodes that descend from the same parent.

As an example of the impact of evolutionary model and sampling scheme on the observable consequences of the restrictions encoded in the graphs, *Supplementary Material Section* 15 shows inferred oncogenetic trees for each of the three true graphs without conjunctions (trees), under several scenarios, including inferred oncogenetic trees with a sample size of 26000 (to minimize sampling variation). The inferred trees are perfect, or almost perfect, reconstructions of the topology of the true tree, but the estimates of the conditional probabilities show large differences. The variations are in directions we would expect both between S.Time and among models, as well as among nodes. Even if the above results are intuitively reasonable, they highlight that whereas the topologies of the graphs (the partial orders) encode constraints in the order of mutations, the conditional probabilities (or *λ*s) we estimate and, most importantly, the patterns of co-occurrence of mutations and the sets of clones we observe, will depend crucially on the evolutionary model and sampling scheme. Since the topology reconstructions depend on the patterns observed, it follows that our inferences will be strongly impacted by evolutionary model (and sampling scheme), as we have seen repeatedly in the results. Moreover, examining the consequences of sampling scheme (S.Time and S.Type) and the detrimental effects of having to separate drivers from passengers on the quality of our inferences can only be meaningfully considered with respect to an evolutionary tumor progression model that generates the data.

As mentioned in the Introduction, the above interaction between order restrictions and evolutionary model, and the unavoidable need to interpret parameters in the context of a given evolutionary model, are coherent with the limitations pointed out by Sprouffske et al. [39]: that oncogenetic tree models (and related models) are not really evolutionary models and do not represent ancestral relationships, but only summarize patterns of co-occurrences of mutations across samples. Virtually all studies of methods for inferring order restrictions are susceptible to this criticism, since they simulate data directly from the generative (but non-evolutionary) OT/CBN/Progression Network model. However, the design I have used here completely overcomes this limitation: I have simulated the data using plausible evolutionary models that incorporate the restrictions in the order of mutations via a straightforward effect on the fitness of clones. Moreover, deviations from monotonicity are not added to the model just as an unexplained error term, but are an integral part of the evolutionary model that can be related, for instance, to the genetic context-dependence of the driver/passenger status.

Sprouffske et al. [39] conclude that cross-sectional data can be misleading if we try to infer the order of mutations. But this conclusion is based on a design where a single OT is inferred from a cross-sectional sample where mutations are not restricted to obey a pre-specified set of restrictions. Thus, it is not surprising that the OT fit does not do well. The results of [39] of course highlight that if different subjects have different sets of order restrictions, then no single OT will capture these patterns, a limitation that is already recognized in the early literature on ongenetic trees [14, 15], and that has prompted the development of mixtures of oncogenetic trees [24, 25, 75]. But, by themselves, their results do not show that OTs (or CBNs or DiPs) from cross-sectional data cannot fare well if there is a true underlying set of restrictions that can be represented as a single graph. Quite to the contrary, I have shown here, embedding the restrictions in evolutionary models, that they can do very well and even recover the exact underlying graph (at least under certain scenarios). Moreover, [39] do not show that any particular within-subject method is actually capable of recovering the true paths from their data (they sidestep that problem altogether).

The two key remaining questions to be answered regarding the usage of cross-sectional data, then, are two: 1) whether the accumulation of mutations in cancer progression can be reasonably represented by a single graph that encapsulates restrictions; 2) if 1) does not hold, whether cross-sectional methods such as mixtures of oncogenetic trees can recover the set of different restrictions. If the answer to 1) is positive, the results of this paper indicate that we have methods that can recover those relationships, and these results also highlight possible avenues to improve them. But question 1) is one that neither this study nor the one of [39] can answer (I simulated data assuming a Yes to that question and [39] assuming a No). Question 2) remains to be thoroughly addressed, and neither this paper nor [39] shed light on the matter. If the answer is negative, then we need to start focusing on within-individual data, which are much harder to obtain.

Nevertheless, the approaches in this paper provide a principled and general way to address that question by simulating data under scenarios were there is no single set of restrictions in common to all subjects, and examining the consequences both for our methods of inferring trajectories and for the data patterns themselves (so as to try to infer, from them, whether or not there is a single set of restrictions).

## Acknowledgments

C. Lazaro-Perea, L. del Peso and J. Poyatos for comments on the ms. C. Lazaro-Perea for help with diagrams and tables. I. B. Diaz for rooting many graphs. R. Salom´on for discussion about CBNs and OTs. H. S. Farahani, T. Graham and R. Datta, M. Gerstung, K. Hainke, C. D. McFarland, W. Mather and L. Tsimring, T. Sakoparnig, and K. Sprouffske for answering questions about their papers and methods and/or providing code of the implementations of their algorithms. Partially supported by Project BIO2009-12458 from the Spanish MINECO.

